# Synchrotron FTIR as a tool for studying populations and individual living cells of green algae

**DOI:** 10.1101/808220

**Authors:** Kira L. Goff, Thomas Ellis, Kenneth E. Wilson

**Affiliations:** University of Waterloo; University of Saskatchewan

## Abstract

Fourier transform infrared (FTIR) spectromicroscopy was used to study individual living cells of three closely-related species of the green algae *Chlamydomonas*. This study differentiated these three species based on differences in lipid and protein profiles, as well explored sources of variation in our measurements. Significant spectral variation was observed between individual cells within a single culture, as well as between control cultures of a species obtained on different days. Despite this, we were able to differentiate between the three close-related species. Differences in the spectra were confirmed using principal component analysis. Understanding the interplay of underlying variation and the degree of induced spectral differences is essential for the deployment of FTIR measurements in both bulk cultures and for individual living cells.

## Introduction

Fourier transform infrared (FTIR) spectromicroscopy offers a unique opportunity to study individual living cells. It allows simultaneous observation of differences in classes of biomolecules of interest in their natural environment and native state. Spectroscopic observations of proteins and lipids provide the opportunity to study functional impacts of changes in cellular components or metabolism. It is a non-destructive technique that allows for simultaneous assessment of all biomolecules. Traditional work involving the extraction of individual cellular components such as lipids, proteins, or DNA allows for in-depth bulk analysis of one class of compound. However, extraction processes kill cells and homogenize the compounds in question across large numbers of cells in order to obtain sufficient sample sizes. Extraction also removes other compounds of interest, limiting the ability to observe interactions between different cellular components. While FTIR does not allow us to isolate individual biomolecules, previous studies have identified classes of biomolecules such as lipids and proteins, and mapped their intracellular distribution ^1,2^, monitored conformational changes over time ^3–5^, accumulation or depletion of metabolic products ^1,6–8^, and assessed cellular responses to stress or the cell cycle ^5,9–11^.

FTIR spectromicroscopy is an emerging technique when applied in the field of biology. It has seen some utilization in phycological studies, where it has been used to study *in vivo* algal metabolism and fundamental differences in physical structure ^6,7,11–14^. Previous work with individual living cells of *C. reinhardtii* observed the production of ethanol due to fermentative anaerobic metabolism ^7^ and demonstrated the effects of oil sands naphthenic acids on the secondary structure of cell wall proteins ^15^.

In this study, we differentiate between closely related species based on their lipid and protein profiles. In attempting to differentiate between three species of the genus *Chlamydmonas*, this study focused on measurement and understanding sources of variation. In doing so, we are studying individual cells of the three species to better assess the challenges and opportunities of FTIR measurement of individual living algal cells.

## Experimental

### Sample Preparation and Equipment

Wild-type *C. reinhardtii* (WT CC-125) was obtained from the Chlamydomonas Resource Center (University of Minnesota). The two environmentally isolated *Chlamydomonas* species, *Chlamydomonas* DJX-H (DJX-H) and *Chlamydomonas* DJX-J (DJX-J), were isolated from a decommissioned pit mine in northern Saskatchewan ^16^. Cultures of *Chlamydomonas* cells were maintained on Tris-Acetate-Phosphate medium (TAP) supplemented with 50 mM arginine and 1.5% agar ^17^. For experiments, cells were transferred to TAP liquid media in sterilized Erlenmeyer flasks, and placed on an orbital shaker at 120 rpm under continuous, uniform light conditions of 100 μmol photons m^−2^s^−1^.

FTIR measurements using individual algal cells were carried out at the Canadian Light Source synchrotron facility (Saskatoon, SK) on the Mid-Infrared beamline (MidIR, 01B1-01) using a Bruker IFS 66v/S interferometer coupled to a Hyperion 2000 FT-IR microscope in synchrotron-light transmission mode. For imaging, cells were prepared as per Goff et al. ^7^. Cells in TAP media were centrifuged and 2 μL of cell pellet was resuspended in a solution of heavy water (D_2_O) containing 0.1% agarose. D_2_O was used to shift solvent absorption away from the amide absorbance region to allow clearer examination of protein secondary structure. A low concentration of agarose was utilized to immobilize *Chlamydomonas* cells with minimal impact on cells or their spectra. The cell-heavy water solution was loaded into a custom sample holder designed for the study of living cells. Optical windows were 1 mm BaF_2_ separated by a 15 μm spacer.

Cell size along the long axis for *C. reinhardtii* and DJX-J cells averaged 7-9 μm, and the larger DJX-H cells 12-15 μm. To ensure the selection and isolation of individual living cells, the bottom, set-diameter confocal apertures were selected be to 2-5 μm greater than the diameter of individual cells, which is near the diffraction limit for IR wavelengths in the fingerprint region. Some cells exhibited minor degrees of motion despite the presence of agarose in the media, and this increase in aperture diameter allowed them to remain within the apertures during measurement. The top apertures were adjustable, square, glass apertures, and set to 35 μm. In comparison to using set-diameter top and bottom apertures, this combination allowed for a wider field of view while scanning for new cells and appeared to reduce baseline variation. In comparison to using adjustable square apertures on the top and bottom, the bottom round aperture allowed for better exclusion of media in the measurement spot around the oval to round cells, and improved repeatability between days and samples. Background measurements (1024 scans, resolution of 4 cm^−1^) were taken of the media, in an area distant from any cells. Once an individual cell was isolated, 512 scans were taken (4000-800 cm^−1^) at a resolution of 4 cm^−1^, a process that took approximately five minutes.

### Data Analysis

The raw data obtained from the FTIR scans of individual cells or cell-free media were analyzed using OPUS 7.2 spectroscopy software (Bruker, United States). Several spectral ranges in the mid-IR range were dominated by solvent components: 3200-3700 cm^−1^ (νH_2_O, νHOD), 2200-2700 cm^−1^ (νD_2_O), 1240-1150 cm^−1^ (δD_2_O). Based on the complex biochemical makeup of the living cells, two distinct spectral regions were used for data analysis: the fingerprint region at 1800-900 cm^−1^ and the CH stretching region at 3050-2800 cm^−1^. The fingerprint region was primarily examined in the 1800-1240 cm^−1^ region due to the limiting nature of the overlapping peaks below this point. Spectra were min-max normalized to the amide I peak centred around 1650 cm^−1^. The lowest absorbance unit point in the local baseline between 1800-1600 cm^−1^ was set to zero while the maximum absorption was set to ‘2’ and a linear multiplicative scalar was applied. The CH region was further offset corrected to set the local baseline at 3040-3010 cm^−1^ to ‘0’. No further baseline correction was done. When 2^nd^ derivative spectra were generated, seven smoothing points were used.

Principal component analysis (PCA), an unsupervised statistical approach that extracts and displays important variability, was performed using The Unscrambler X (CAMO Software, Olso, Norway). Normalized spectra were imported from OPUS, and the 1800-1240 and 3050-2800 cm^−1^ regions were analyzed separately. Data preprocessing for the 1800-1240 cm^−1^ region consisted of the application of an extended multiplicative scattering correction (EMSC) to correct for additive, multiplicative, and wavelength-depended spectral effects. This corrected for factors such as differences in path length (different thicknesses in the 15 μm sample space between sample holder loadings), scattering caused by slight differences in the angle of the paired windows inside the sample holder, interference patterns from alignment of the optical windows, condenser and microscope apertures, Mie scattering, and similar factors. Loadings for the principal components that define PCA space are displayed in figures as horizontal lines that span the spectral regions that drive the principle components. EMSC-corrected spectra were visually compared to offset-corrected spectra to ascertain that the spectral changes were relatively minor. Initial analysis of PCA of the CH stretching region showed that while the spectra of different treatments separated out nearly perfectly in PCA space, the loadings were representative of baseline slopes and associated overall intensity differences, rather than changes in the shape, relative size, and position of peaks. To allow for more meaningful comparisons in the CH stretching region, the EMSC correction was followed by the generation of 2^nd^ derivative spectra (Savitzky-Golay, 7 smoothing points) which were used for PCA analysis. Once a PCA was generated, loadings and correlational loadings were examined to see which spectral ranges influenced each principal component axis.

## Results

### Intercellular variation

To minimize sample variation, cell cultures were grown at the same time and cells from those cultures were measured on the same day. When examining the FTIR spectra of individual living cells obtained from the same culture on the same day, three types of variation were observed. The types of observed variation were: 1) Variation around a mean (Figure 1); 2) Individual outliers (Figure 2); 3) Presence of subpopulations of cells (Figure 3). Despite being haploid clones, the cells in the culture were not all identical when measured using the FTIR spectromicroscopy system (Figure 1). Even following min-max normalization to the Amide I band, differences in absorption were observed (Figure 1A). When measurements were taken of 10 different cells ((Figure 1A) no two spectra were identical to each other. In addition, three of these cells (F, G, and I) are markedly different from the others in at least one particular peak or region of their spectra. Cell F (Figure 1A, green line) exhibits a higher baseline in the 1500-1240 cm^−1^ region, cell G (Figure 1A, dark blue line) has a stronger 1740 cm^−1^ peak, and cell I (Figure 1A, purple line) exhibits a much stronger 1080 cm^−1^ peak. Comparing the averages generated by all ten of these cells (Figure 1B, blue) versus the average generated by removing these outliers (Figure 1B, orange), it can be observed that the influence of cell to cell variations around the mean were minimal when considering an averaged spectrum.

**Figure 1.**
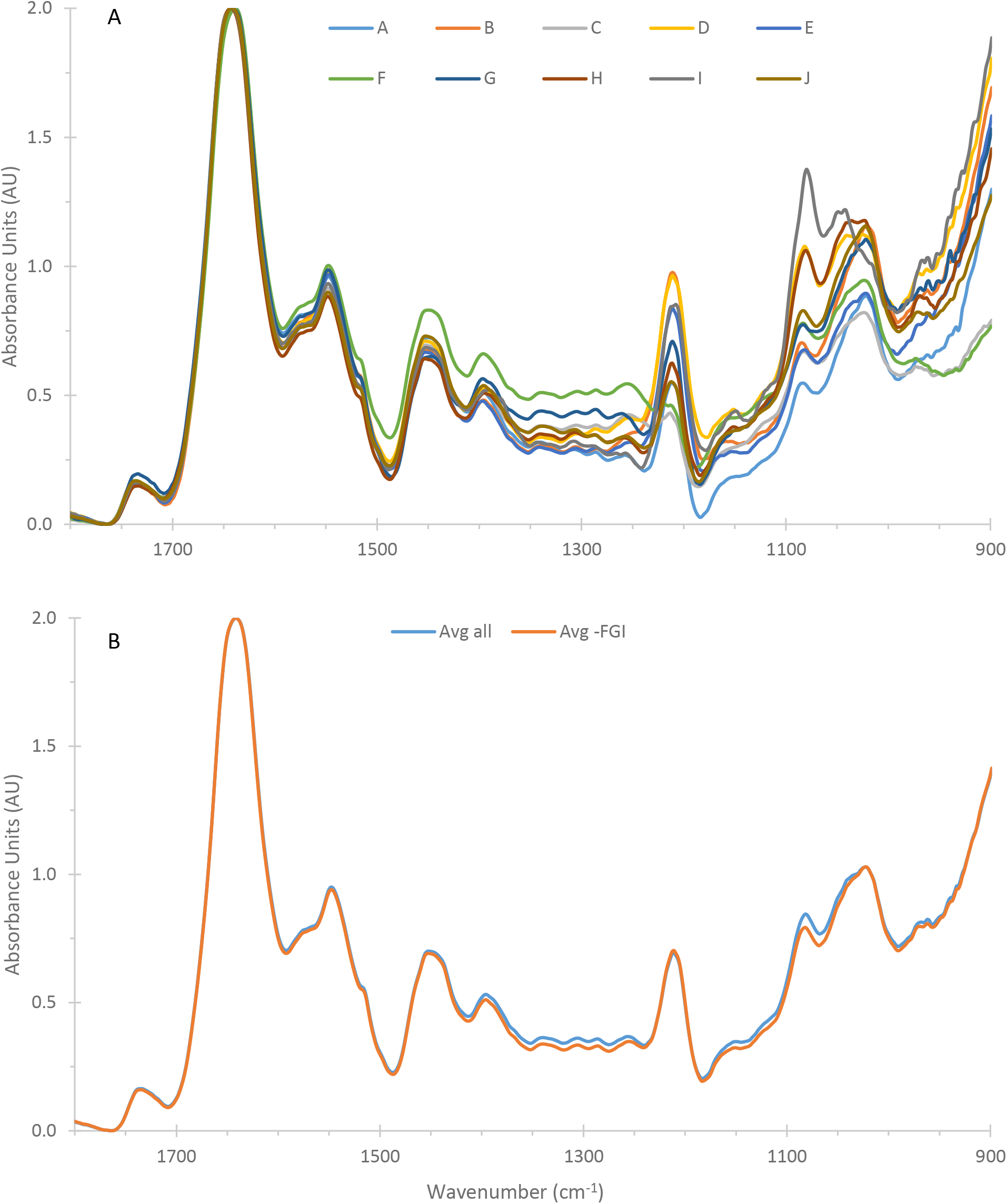
A) Spectra of 10 individual cells of DJX-J, including three cells (F, G, I) that differ from the other spectra in one region. B) Spectra generated by averaging all 10 cells (blue), or 7 cells, removing FGI, each of which have one abnormal peak (orange), showing the minimal influence on the resultant average.

**Figure 2.**
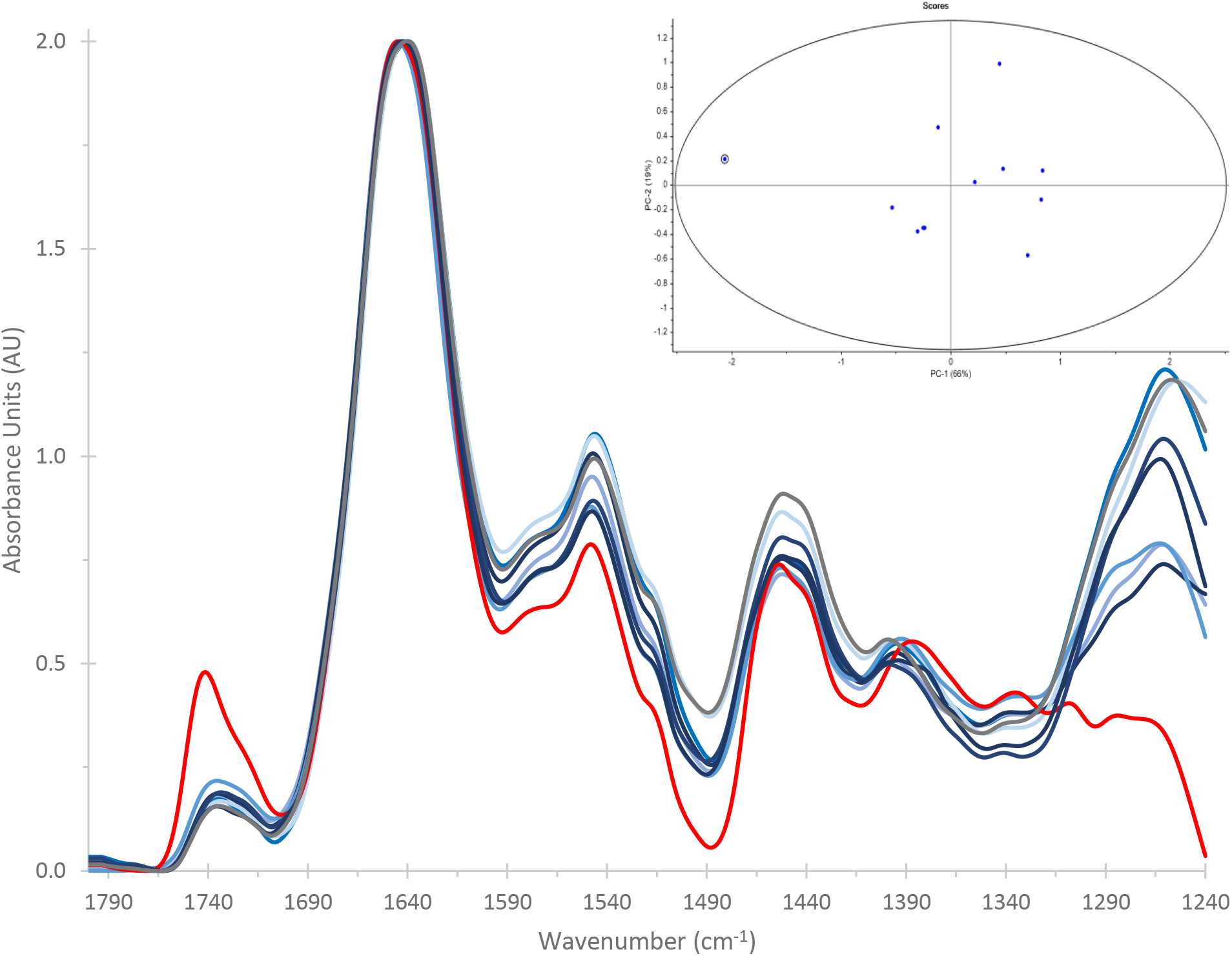
Spectra of individual living cells of Chlamydomonas DJX-H obtained from a single culture. Spectra in blue exhibit inherent intercultural variation in line with the minor, continuous variation expected from cells exhibiting slight differences in physiology, metabolism, and biochemical composition. The red spectrum is an outlier identified by strong, discrete differences. These are most obvious in the ~1740 cm^−1^ lipid carbonyl, 1398 cm^−1^, 1308 cm^−1^, and 1260 cm^−1^ peaks. Inset: PCA confirming outlier status.

**Figure 3.**
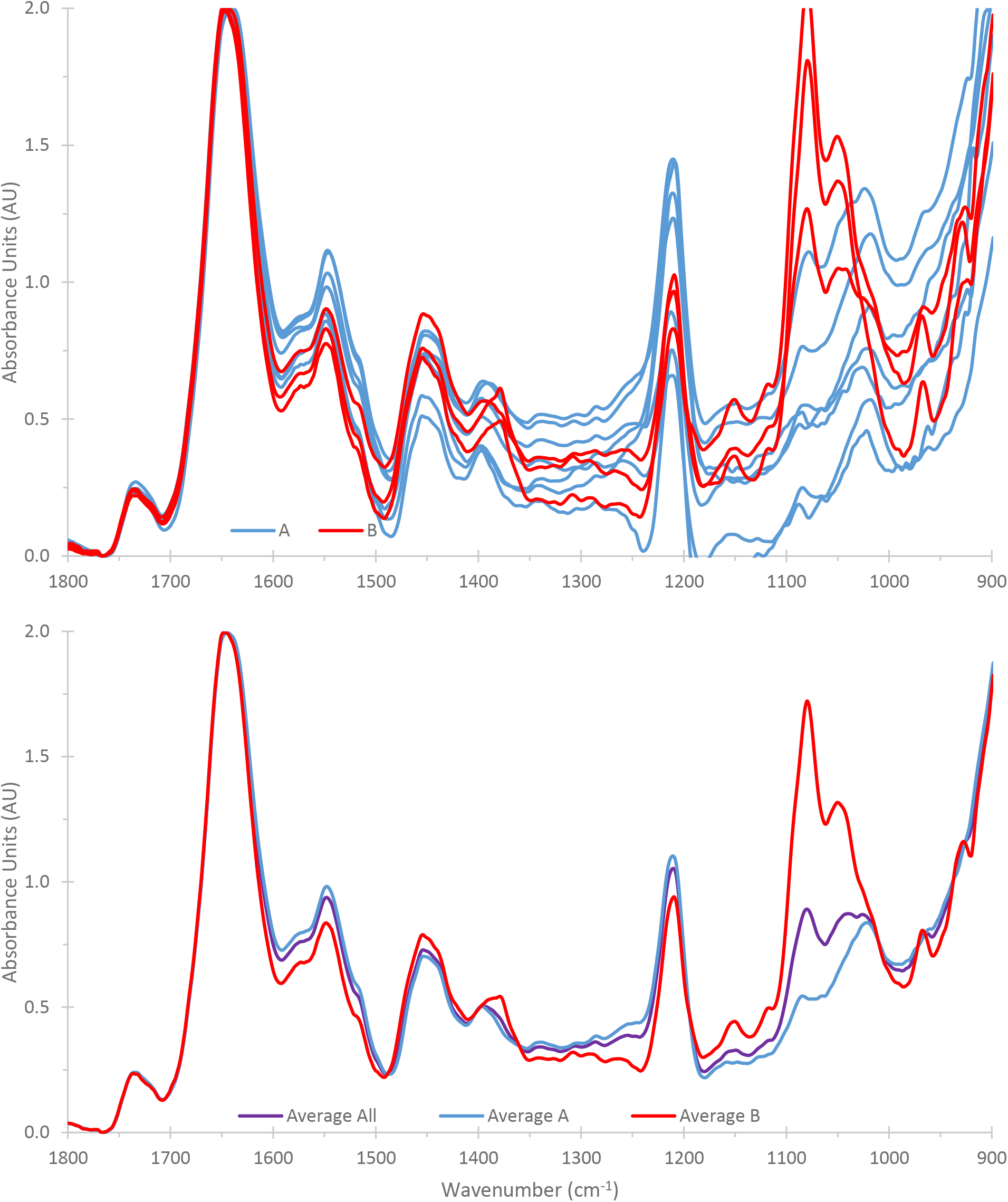
A) Identification of two subpopulations of cells within a culture of Chlamydomonas DJX-J. Subpopulation ‘B’ is differentiated by the strength and position of 1398 cm^−1^, as well as the strength of 1308 cm^−1^, 1080 cm^−1^, 1040 cm^−1^, and 1020 cm^−1^. B) Influence of the presence of two subpopulations of cells on the average procedure.

When considering spectra from individual cells, there were rare occasions when a single, extremely different spectrum was observed (Figure 2, red line). The spectrum from this particular cell exhibited differences in the strength and shape of the 1740 cm^−1^ lipid carbonyl peak and the loss of the strong peak centred around 1260 cm^−1^. There were also differences are present in the amide I:amide II ratio, and reshaping of the 1390 cm^−1^region (Figure 2). One way to assess if the overall spectrum is different from those of other cells measured from that sample is to perform PCA (Figure 2 inset). By comparing the spectrum of the suspected outlier using PCA, one can set a priori limits on what represents an outlier. This allows the researchers to limit noise due to artifacts, or dead cells, for example, without selection bias. The spectrum noted as an outlier in Figure 2 clearly separates from the rest of the spectra in two-dimensional PCA space (inset). This dovetails with visual observations of spectral differences that are distinct rather than continuous (Figure 1) or part of a subset of cells (Figure 3). This is a spectrum that would be removed from the data pool before further analysis occurred because of the out-sized influence this spectrum would have on any averages generated.

An additional complication can arise if, rather than a single outlier, there are multiple populations of cells. An example of this situation was observed within a DJX-J culture (Figure 3). In this case, two populations were delineated by differences in: absorption in the amide II band, the strength and position of the 1398 cm^−1^, and the strength of the 1308 cm^−1^, 1080 cm^−1^, 1040 cm^−1^, and 1020 cm^−1^ peaks (Figure 3A population 1, blue; population 2, red). The existence of multiple cells with such different spectra can exert a large difference on overall spectrum averages (Figure 3B, blue, population A; red, population B; purple, all cells). This contrasts significantly to the situations of individual variation around the mean (Figure 1B) and a single outlier (Figure 2). Combining the spectra from cellular subpopulations has a very strong influence on the average.

### Differentiating between closely related species

These data handling considerations were applied in the construction of Figures 4-6. For these figures, the spectra presented are the average of data collection across four cultures on four days (*C. reinhardtii*), or five cultures on five days (DJX-H, DJX-J). Each point in the PCAs inset into the B) figure represents the average of 7-9 spectra that passed the above criteria on a given sampling day. The spectra displayed for a species are the average of averages. While all of these figures show some extent of day to day variation for each species, PCA confirms that there are real and repeatable spectral differences observed between species.

**Figure 4.**
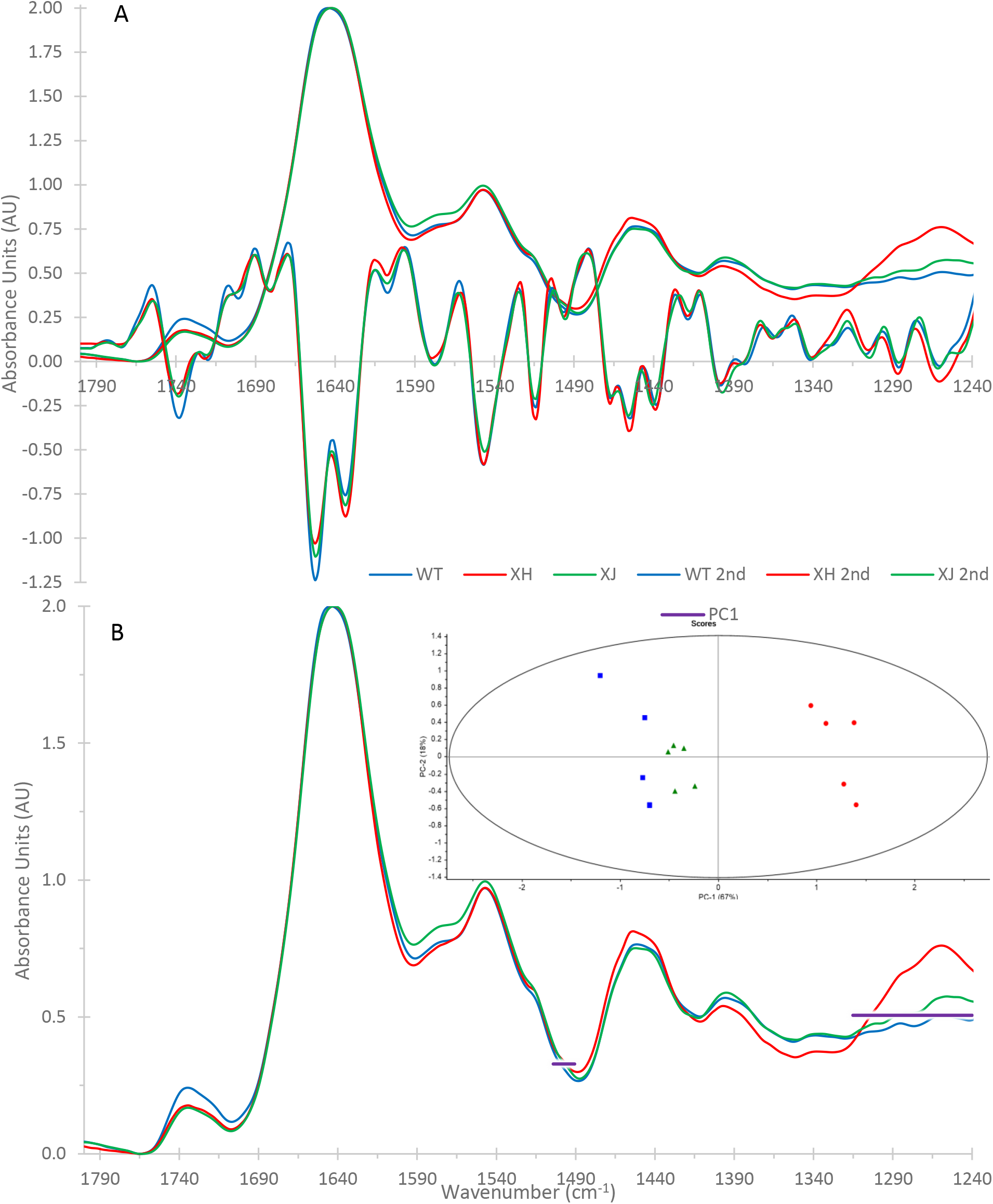
FTIR measurements of the 1800-1240 cm^−1^ region in Chlamydomonas reinhardtii (blue), Chlamydomonas DJX-H (red) and Chlamydomonas DJX-J (green). Spectra are the averages of the individual spectra used in principal component analysis and have been normalized to the amide I peak. A) Average spectra and scaled 2^nd^ derivatives; B) Average spectra overlain with the loadings generating inset image (PC1-3: 67%, 18%, 5%).

We anticipate that due to variation in the lipids and proteins found in different species of the genus *Chlamydmonas*, it will be possible to differentiate between cells of different species based on the spectra we obtain using synchrotron-based FTIR spectromicroscopy. To focus on the absorbance bands related to proteins and lipids, close examination of the 1800-1240 cm^−1^ and 3050 – 2800 cm^−1^ regions was performed. Beginning with the 1800 to 1240 cm^−1^ region (Figure 4), the most immediately obvious difference between these three species was the presence of a strong, broad peak centred around 1260 cm^−1^ in the spectra of DJX-H cells (Figure 4, red). This peak started around 1310 cm^−1^ and ran through the cutoff at 1240 cm^−1^. The intensity of the 1260 cm^−1^ peak observed in spectra from DJX-H cells is such that the use of principal component analysis to differentiate between these three species of *Chlamydomonas* (Figure 4 B) results in loadings that are almost entirely driven by the 1260 cm^−1^ peak alone (Figure 4 B, inset, loading represented by horizontal line through the spectral regions associated with the principal component). Data analysis after exclusion of this peak (using 1800-1310 cm^−1^, Figure 5) allows for better separation of *C. reinhardtii* (blue) from DJX-H (red) and DJX-J (green) across the spectrum. The 1260 cm^−1^ region is associated with PO_4_- and the amide III protein peak 18, and as such has multiple overlapping biological assignments including proteins, phospholipids, nuclear and chloroplastic DNA and RNA, and phosphorylated storage products or photosynthetic proteins ^19,20^. One strong possibility is polyphosphate compounds, which in their pure form absorb primarily through this region and below ^21^. No firm assignment can be made based on available data.

**Figure 5.**
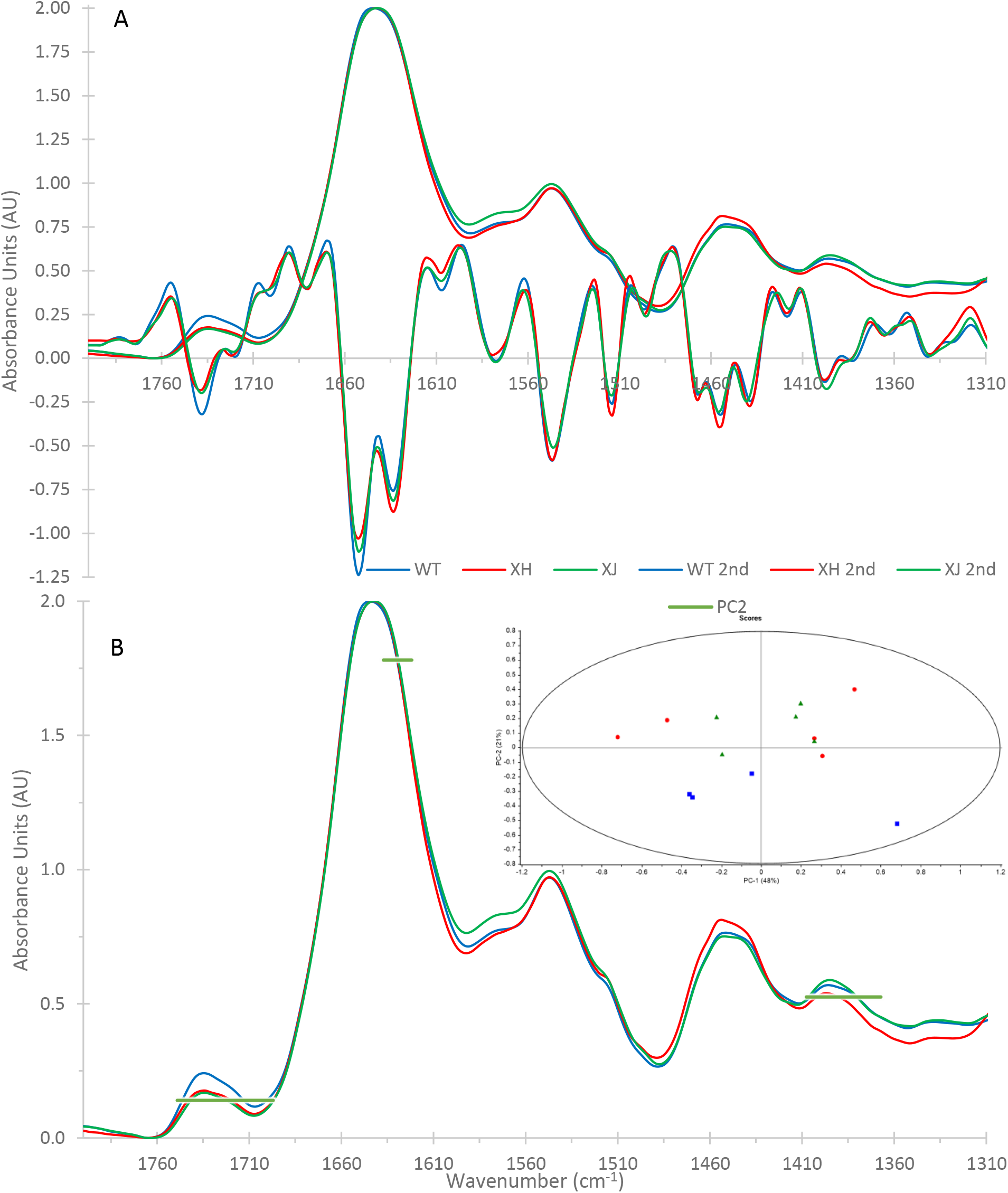
FTIR measurements of the 1800-1310 cm^−1^ region in Chlamydomonas reinhardtii (blue), Chlamydomonas DJX-H (red) and Chlamydomonas DJX-J (green). Spectra are the averages of the individual spectra used in principal component analysis and have been normalized to the amide I peak. A) Average spectra and scaled 2^nd^ derivatives; B) Average spectra overlain with the loadings generating inset image (PC1-2: 48%, 21%).

Beginning at the left-hand side of Figure 5B, one assignable difference between the spectra obtained from cells of these three species is the shape and intensity of the ~1740 cm^−1^ lipid carbonyl peak (Figure 5). The second derivative of this peak contains three primary components with minimums at 1740 cm^−1^ (esters), 1720 cm^−1^ (carboxylates), and 1700 cm^−1^ (unassigned), as per Movasaghi et al. ^18^. The ~1740 cm^−1^ lipid carbonyl band differed in both shape and intensity when comparing spectra from *C. reinhardtii* (blue), and DJX-H, and DJX-J cells (red, green). Spectra from cells of DJX-H and DJX-J exhibited a lower intensity of the 1740 cm^−1^ and the 1700 cm^−1^ components.

Comparing the spectra revealed differences in protein secondary structure between these three species. Second derivative components of the amide I peak are indicative of the secondary structure of the proteins being measured ^22–24^. The peak position of *C. reinhardtii* (1644 cm^−1^) was higher when compared that of DJX-H and -J (1642 cm^−1^); as a result the Amide I peak of *C. reinhardtii* is less symmetrical; these differences are reflected in the loadings of PC-2. Examination of the second derivative spectra of these three species revealed a relative enrichment of β-sheets (1633 cm^−1^) in comparison to α-helixes (1652 cm^−1^) in DJX-H and DJX-J when compared to *C. reinhardtii*. This is the main separator of DJX-H and J, from *C. reinhardtii* in the PCA (Fig. 5B inset) and is reflected in the PCA loadings. Changes in the shape of the amide II peak were more subtle and appeared to be related to slight increases in the width of the 1547 cm^−1^ and 1515 cm^−1^ components, which when spectra from *C. reinhardtii* cells were compared were broader in DJX-J and narrower in DJX-H cells (Figure 5A).

Other differences are present in the fingerprint region below the amide II peak in spectra obtained from cells of the three species. This region becomes increasingly complicated due to numerous overlapping absorbance peaks. The shape of the ~1450 peak is slightly different when comparing spectra from each of the three species (Figure 5). Looking at the second derivative one sees relative contributions from three distinct components (1467 cm^−1^, 1456 cm^−1^, and 1440-1438 cm^−1^). These second derivative components are respectively associated with δCH_2_ and δCH_3_ of lipids and proteins ^18^, δHOD absorption ^25^, and HOD exchange off proteins associated with the amide II peak ^23^. The spectra of DJX-H cells show an overall greater intensity in the 1450 cm^−1^ peak, as well a stronger relative contribution from the second derivative 1456 cm^−1^ component (δHOD absorption) (Figure 5A). Differences in peak shapes and intensities were present in the 1398 cm^−1^ and 1340 cm^−1^ peaks, but they are relatively subtle, and the peaks were quite broad. These peaks each have two variable, second derivative components with several possible assignments, including protein, lipids, and carbohydrates ^18^.

Principal component analysis of the 1800-1310 cm^−1^ region of the spectra obtained from these three species (Figure 5B insert) found discrete areas of differences between their spectra, and therefore allowed for the separation of the environmental DJX species from the laboratory species *C. reinhardtii*. This was based on PC-2, which was driven by differences in the 1740 cm^−1^ lipid peak, the relative enrichment of β-sheets in the DJX species, and differences in the width of the 1398 cm^−1^ peak.

The high wavenumber CH stretching region (3050-2800 cm^−1^, Figure 6) revealed a variety of differences between *C. reinhardtii* and the DJX-species. The overall higher intensity of the CH_2_ and CH_3_ peaks in *C. reinhardtii* is consistent with this species’ correspondingly stronger 1740 cm^−1^ peak, suggesting a greater lipid accumulation (Figure 5). The spectra of cells of *C. reinhardtii* (blue) also exhibited a more intense 3010 cm^−1^ peak assigned to C-H functional groups. Other spectral differences of interest between these cells were found in the position and width of the CH_2_ and CH_3_ peaks. In spectra obtained from DJX-H and DJX-J cells, CH peaks exhibited higher peak positions and greater peak widths when compared to *C. reinhardtii*. This suggests a greater diversity of lipids present in these cells compared to *C. reinhardtii*. The DJX-species also showed different absorptions around the unassigned CH second derivative component at 2995 cm^−1^. The differences noted above reflect the abundance and diversity of lipids, leading to a PCA separation between the DJX species and *C. reinhardtii* (Figure 6 B, inset).

**Figure 6.**
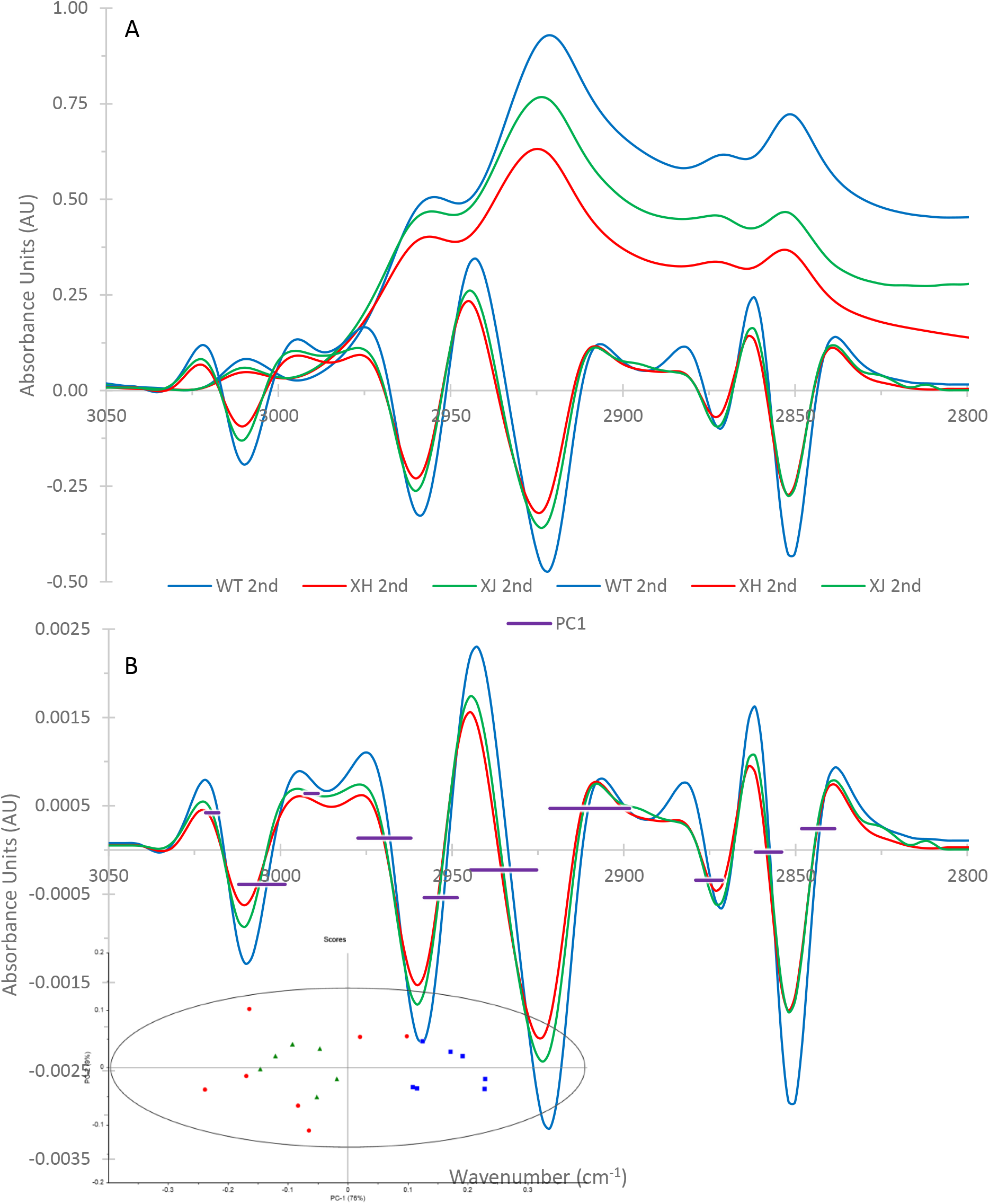
FTIR measurements of the 3050-2800 cm^−1^ region in Chlamydomonas reinhardtii(blue), Chlamydomonas DJX-H (red) and Chlamydomonas DJX-J (green). Spectra are the averages of the individual spectra used in principal component analysis and have been normalized to the amide I peak, then offset corrected 3050 – 3000 cm^−1^. A) Average spectra and scaled 2^nd^ derivatives; B) Average of 2^nd^ derivative spectra overlain with the loadings generating inset image. (PC1-3: 57%, 18%, 7%).

### Influence of intercultural variability on experimental design

Figures 4-6 demonstrate that the variation observed between cells in different cultures (Figures 1-3) still allows for discrimination between closely related species. However, the degree of variation present within and between cultures can still be a confounding factor for data analysis. The influence of day-to-day variation on species differentiation can be seen in the PCA insets of Figures 4B, 5B, and 6B. In this instance, the spectral differences between these closely related species were such that we were able to differentiate between them, despite this variation. Depending on how subtle spectral difference of interest are – between species, or in response to external stimulus – this might not always be the case.

The extent to which day to day variation acts as a confounding variable that might obscure desired observations will depend upon how clear or subtle cellular responses are to a given stimulus. In some cases, this degree of variation may almost entirely obscure experimentally-induced spectral variation. Figure 7 demonstrates the influence of variation between spectra obtained from different cultures on different days (day 1, blue; day 2, red; each point represents the measurement from an individual living cell). Spectra from each day include both control spectra and the spectra of cells exposed to oxidative stressors, which produce relatively subtle spectral effects. Even with stressor exposure, the two cultures are completely separated in PCA space. As a result, comparison of control and treatment cells obtained on different days would inherently conflate the spectral influence of experimental treatments and intercultural variability.

**Figure 7.**
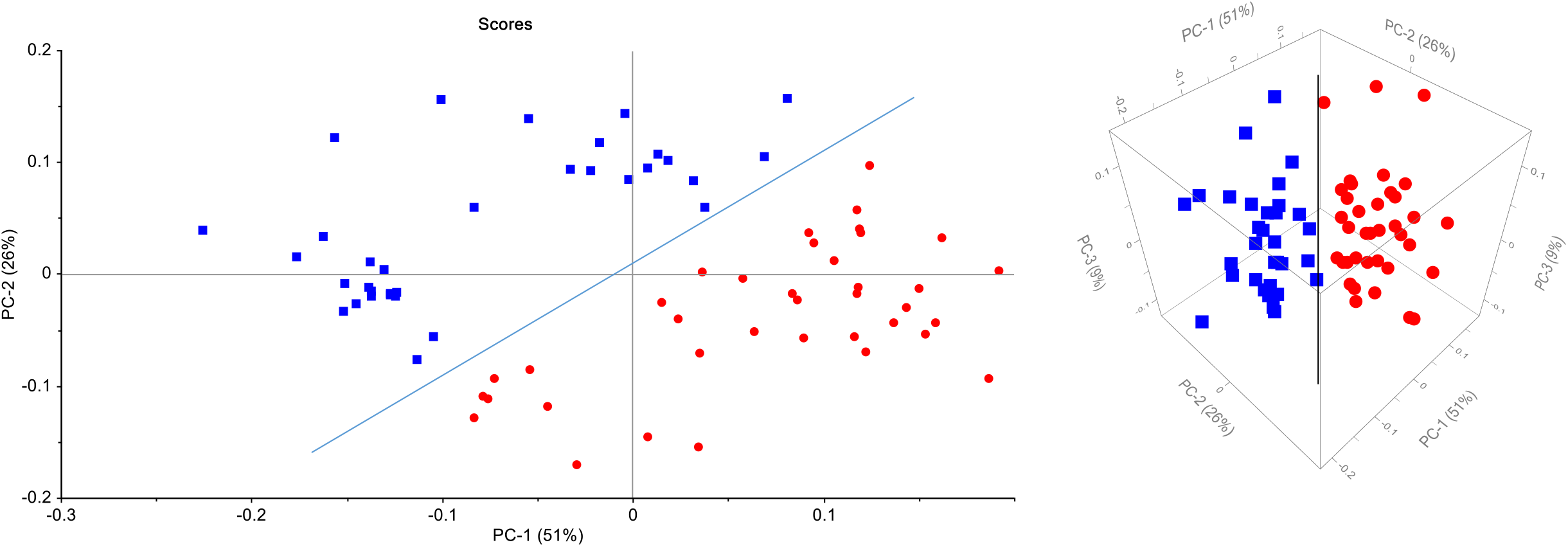
Demonstration of spectral separation in PCA space by day/culture for Chlamydomonas DJX-H. Blue spectra obtained day one, red obtained day two. Diagonal line added for reference.

## Discussion

FTIR spectromicroscopy is a very powerful and sensitive tool with a broad range of potential applications for the study of individual living cells. However, one must consider that the sensitivity of the technique and the difficulty inherent in working with living cells results in both challenges and opportunities.

The individual living cells measured in this work are haploid and clonal and therefore theoretically exhibit minimal variation. However, due to differences in cell age, culture maturity, and even physiological differences, noticeable amounts of spectral variation were observed. The three types of variation highlighted in Figures 1-3 (variation around the mean, individual outliers, or the presence of multiple subpopulations of cells) each carry different implications for data handling and processing. Variation around the mean, as shown in Figure 1, yields spectra that can be safely combined without outsized influence on the resulting average. This is particularly important to know when working with the small sample sizes caused by limited material and availability of synchrotron beam line time. One needs to optimize their measurements to get the most reproducible data in the shortest time possible. However, some spectra obtained from individual cells clearly do not represent the average cells in a population. This was highlighted as the case in the presence of extreme outliers (Figure 2) and subpopulations (Figure 3). These non-representative spectra present greater challenges for data analysis and require a greater degree of user oversight.

In the case of the outlier spectra (Figure 2), the absorbance differences not only had a strong influence on the calculated average but was indicative of significant alterations in the physiology and biochemistry of that cell. Observed differences in the strength and shape of the 1740 cm^−1^ lipid carbonyl peak, the altered amide I:amide II ratio, reshaped 1390 cm^−1^, and the loss of the strong peak centred around 1260 cm^−1^ suggested major differences in lipids, proteins, and phosphates. If true, this cell was not representative of the bulk of the culture. Of course, the changes could be due to measuring artifacts such as movement of the cell in the measurement area. However, it is unlikely that the cell moved during measurement to produce these results, because cell location in the measuring area was double checked at the end of each observation. The movement of the cell away from and then back towards the measurement spot is extremely unlikely, and cells that did exhibit motion generally yielded noisy, low-intensity spectra, unlike the outlier spectra presented in Figure 2. As such, it is likely that this cell is chemically different than the others in the culture, be it related to cell age, health, reproductive status, or other unknown factors.

Strong outliers were differentiated from cellular subpopulations (Figure 3) based on the occurrence of multiple cells exhibiting similar spectral features. In this case, subpopulations of cells exhibiting the same features were found across multiple cells lines on multiple days. In multiple experimental runs, cultures were identified with a small subpopulation of cells defined by increases at 1477 cm^−1^ (w), 1412 cm^−1^ (sh), 1377 cm^−1^ (m), 1308 cm^−1^ (w), 1152 cm^−1^ (m), 1118 cm^−1^ (sh), 1080 cm^−1^ (vs), 1051 cm^−1^ (s), 1020 cm^−1^ (sh), 969 cm^−1^ (m), and 931 cm^−1^ (m). These peaks have overlapping assignments, but can be generalized to proteins and lipids (δCH_2_, δCH_3_, COO-: 1477, 1398, 1377, and 1308 cm^−1^), and phosphorous containing compounds such as phospholipids, DNA, RNA, and sugar-phosphates (1152, 1118, 1080, 1050, 1020, 969, 931, and 889 cm^−1^) ^18^. Overall analysis and the strength of the changes in the phosphate region suggest that these differences are more likely due to changes in DNA/RNA, phospholipids, or phosphorylation of sugars or proteins. This specificity suggests the possibility that the variance being captured here is reflective of cells in different portion of the cell cycle, after DNA synthesis has begun but prior to cell division being initiated.

The presence of these types of variation - variation around the mean, outliers, and subpopulations - strongly supports the utilization of individual living cells for study when possible. Bulk or averaged measurements are only representative of variation around the mean (Figure 1). In the case of outliers or subpopulations, any averages generated will ignore the complexity of the system and may result in skewed comparisons going forward. The use of bulk culture measurements is predicated upon the assumption that cellular populations are distributed predictably about a mean, and we have shown that it often fallacious. If cultures or cellular populations contain strong outliers or distinct subpopulations, it is reasonable to question whether this alters cellular stress response (data in preparation for publication). It is possible that in many cases – depending on the variable being studied – the use of bulk cultures may still be acceptable, but it is not reasonable to proceed with this assumption without ascertaining if this is the case for a particular study system. Dealing with subpopulations of cells is normally accomplished by comparing the subpopulations to spectra obtained on other days and discarding the spectra from the subpopulation that is distinctly different from the overall dataset.

Having a strong grasp of the types of variation present in our cell cultures and the resulting spectra, allowed the investigation of using FTIR spectromicroscopy to differentiate between closely related species of green algae. In this case the model alga *C. reinhardtii* was compared to two *Chlamydmonas* species isolated from a water source in northern Saskatchewan. It was predicted that the algal species would have different biochemical makeups, which would be distinguishable by FTIR at the single cell level.

The most striking feature visible in the fingerprint region of these cells is the strong peak centred around 1260cm^−1^ in the spectra of DJX-H cells. PCA of these spectra confirms its importance in differentiating between these three species, as it dominated the loadings of the fingerprint region. Indeed, differences were found in the overall amounts of lipids present in the cells (lipid abundances) as well as in the relative amounts of various types and classes of lipids (lipid cohorts). One immediately obvious and assignable difference between these three species was found in the shape and intensity of the ~1740 cm^−1^ lipid carbonyl peak (Figure 5). Using PCA, the averaged spectra of the three cell types could be compared and differences used to group cells that are most similar. The 1740 cm^−1^ region was part of PC-2 that allowed differentiation between *C. reinhardtii* and DJX-species. Spectra from cells of *C. reinhardtii* exhibited overall greater intensity in the CH stretching region (Figure 6), consistent with this species’ correspondingly stronger 1740 cm^−1^ peak (Figure 5), suggesting greater lipid abundance with respect to proteins. The differences in the shape of the 1740 cm^−1^ peak and CH stretching regions also suggestions that *C. reinhardtii* carries a different compliment of lipids than the environmental DJX-species.

Beyond differences in the overall intensity of the CH stretching region, important information was gleaned from differences in the peak position and peak widths of the CH_2_ and CH_3_ peaks. The spectra obtained from DJX-H and DJX-J cells exhibited CH peaks at higher positions and greater peak widths when compared to *C. reinhardtii*. These differences, specifically in νCH_2*s*_ peak position (2851 cm^−1^ in *C. reinhardtii*, 2853 cm^−1^ in the DJX-species) and peak widths, indicated increased membrane disorder in the DJX-species ^26^. The DJX-species also exhibited different absorptions around the unassigned CH second derivative component at 2995 cm^−1^. This may be due to interaction of the upshifted νCH_3*s*_ peak and another unidentified contribution at 2885 cm^−1^, where the weaker contributions overlap with the stronger peaks. Just as PCA of the 1800-1310 cm^−1^ region (Figure 5) showed clear separation between *C. reinhardtii* and the DJX-species, PCA of the CH stretching region (Figure 6) showed clear separation between *C. reinhardtii* and the DJX-species. Taken together, this information suggests that the life history of these species (originally isolated from a field in the 1940s, raised under laboratory conditions since then, *C*. *reinhardtii*; uranium mine pit, DJX-) influences not only lipid abundance and cohort but also membrane structure.

It was also possible to differentiate between *C. reinhardtii* and the DJX-species on the basis of protein secondary structure (Figure 5). This was most obvious in examination of the second derivative of the amide I peak. These differences are more difficult to assign to specific physiological differences between species but are consistent and repeatable. The altered prevalence of α-helixes, β-sheets, turns, and globular proteins suggests that there are significant differences in the protein compliments in these cells.

We have shown it is possible to differentiate between closely related green algal species using FTIR spectromicoscopy. PCA analysis was used to highlight these differences. An analysis of the PCA loadings was used to refine the analysis and interpretation of the spectra.

Understanding the differences in various regions of the spectra obtained from individual cells was enhanced by studying the principal component loadings. Appreciating how they drive the distribution of spectra in PCA space is essential to the usage of PCAs. For example, the exclusion of the 1260 cm^−1^ peak from the PCA analysis of the three species allowed for the selection of the region of the spectra that could provide the most informative comparison between spectra from cells of the three species. Similarly, analysis of the PCA plots generated from the CH region yielded clean separation in all cases, but examination of the loadings showed that the driving differences were related to baseline factors and overall spectral absorption. This informed the decision to use second derivatives for analysis in this region, to better represent true spectral differences.

In addition to the variation observed within cultures, significant spectral differences were also observed between control cultures measured on different days (Figure 7). Understanding this underlying variation is essential for experimental design. In some cases, this degree of variation may not impair spectral cross-comparisons, and in others it may almost entirely obscure experimentally-induced spectral variation. Any study that attempts to quantify or qualify spectral changes in response to an exposure or stress must make sure that they have appropriate controls in place. In our case, day-to-day variation between cultures was not sufficient to disrupt out attempts to differentiated between closely related species under control conditions (Figures 4-6). However, when it came to the effects of induced stressors (Figure 7), day-to-day variation was a confounding factor: control cells from one culture could not be used as control spectra for cells from another culture on another day.

Visible differences in peak shape, position, and ratios allowed us to infer that the three algal species contain different relative amounts of compound classes such as lipids and proteins. Differences in the shapes of these peaks also allow the observation that there are interspecies differences in the lipid cohort and protein secondary structures. These differences are expected to reflect the different evolutionary paths that the three species have taken to adapt to survive in their natural environment.

## Conclusions

FTIR spectromicroscopy of individual living cells is a powerful tool when used carefully. The capacity of FTIR to differentiate between species of cells based on their biochemical makeup, is useful only if proper experimental design is grounded in the understanding of the inherent differences present in measurement populations. The study of individual living cells results in greater degrees of complexity in data analysis, but the data obtained is less prone to skew by outliers or the presence of cellular subpopulations, and gives information about culture health and complexity obscured by bulk measurements. Experimental design must take into account both the extent and types of underlying variation and the degree of induced experimental variation. Future studies that non-selectively measure large number of cells can still be viable. However, they should be preceded by study of individual cells to better understand the degree and types of variation present within your population. The degree of inherent variation found within and between cultures, when compared to the intensity of changes induced by experimental conditions, is an essential factor for designing experiments.

Despite these challenges, it was possible to use FTIR to differentiate between closely related species of *Chlamydomonas*. Overall, the spectra obtained from the DJX-species were more similar to each other than to *C. reinhardtii*, the model lab species. These differences in membrane and protein structure may reflect different physiological adjustments to their environment, suggesting the capacity to use FTIR spectromicroscopy to examine how cells respond to environmental stimuli. However, despite their coevolution, DJX-H and -J cells are not identical. The spectra obtained from these two species exhibit reliable spectral differences, particularly the strong, broad 1260 cm^−1^ peak that is only measured in DJX-H cells.

## Acknowledgements

Research described in this paper was performed at the Canadian Light Source, which is supported by the Canada Foundation for Innovation, Natural Sciences and Engineering Research Council of Canada, the University of Saskatchewan, the Government of Saskatchewan, Western Economic Diversification Canada, the National Research Council Canada, and the Canadian Institutes of Health Research. Kira Goff was supported in part by the U of S and NSERC. Kenneth Wilson’s research was supported by a NSERC discovery grant, CFI, NSERC, RTI, and the U of S.

